# Radiotherapy and chemotherapy alter migration of brain cancer cells before cell death

**DOI:** 10.1101/2020.07.23.218636

**Authors:** Michael Merrick, Michael J. Mimlitz, Catherine Weeder, Haris Akhter, Allie Bray, Andrew Walther, Chisom Nwakama, Joe Bamesberger, Honour Djam, Kaamil Abid, Andrew Ekpenyong

## Abstract

Although radiotherapy and most cancer drugs target the proliferation of cancer cells, it is metastasis, the complex process by which cancer cells spread from the primary tumor to other tissues and organs of the body where they form new tumors, that leads to over 90% of all cancer deaths. Thus, there is an urgent need for anti-metastasis strategies alongside chemotherapy and radiotherapy. An important step in the metastatic cascade is migration. It is the first step in metastasis via local invasion. Here we address the question whether ionizing radiation and/or chemotherapy might inadvertently promote metastasis and/or invasiveness by enhancing cell migration. We used a standard laboratory irradiator, Faxitron CellRad, to irradiate both non-cancer (HCN2 neurons) and cancer cells (T98G glioblastoma) with 2 Gy, 10 Gy and 20 Gy of X-rays. Paclitaxel (5 μM) was used for chemotherapy. We then measured the attachment and migration of the cells using an electric cell substrate impedance sensing device. Both the irradiated HCN2 cells and T98G cells showed significantly (p < 0.01) enhanced migration compared to non-irradiated cells, within the first 20 to 40 hours following irradiation with 20 Gy. Our results suggest that cell migration should be a therapeutic target in anti-metastasis/anti-invasion strategies for improved radiotherapy and chemotherapy outcomes.

## 1. Introduction

Although radiotherapy is one of the three most frequent treatment modalities used in the clinic against cancers with longstanding evidence of benefits to patients [1–3], there are surprising reports that ionizing radiation can promote processes that lead to both local tumor recurrence and metastasis such as migration and invasion [4–6]. All three common treatment modalities (radiotherapy, chemotherapy and surgery) target cancer cells. Yet, it is metastasis, the complex process by which cancer cells spread from the primary tumor to other tissues and organs of the body where they form new tumors, that leads to over 90% of all cancer deaths [7–10]. Thus, alongside chemotherapy and radiotherapy, there is a crucial need for anti-metastasis and anti-invasion regimens [11–14]. A critical step in the metastatic cascade is cell migration [15,16]. Cell migration engenders invasion, the primordial step in tumor spreading. Whether radiotherapy and/or chemotherapy promote(s) cancer cell migration or not, strategies that reduce migration are promising anti-metastasis and anti-invasion treatments [12,17] which could in turn improve outcomes for patients. Here, we employ a well-established migration assay to track in real time the migratory ability of both brain cancer cells (glioblastoma) and non-cancer cells (neurons) following irradiation with X-ray photons and paclitaxel treatment. This tracking allows the establishment of time scales involved in radiation-induced alteration of cell migration which can in turn engender the development of effective anti-invasion strategies. Since invasion and metastasis occurs in all cancers [10,18,19] in spite of the very wide variety of cancers with respect to their molecular biology, pathogenesis and prognosis, any contribution to anti-metastatic/anti-invasion treatment strategies would be very significant in the perennial and global fight against cancer.

Furthermore, current clinical practice and research show that radiotherapy is less effective in some cancers than others. For instance, patients of primary brain tumors such as glioblastoma multiforme (GBM) have a median survival of 4-18 months [1,20] with or without radiotherapy. In fact, glioblastomas are among the most radio-resistant, chemo-resistant and aggressive forms of cancer [2,4,20]. The exact mechanisms behind this high migratory ability of GBMs and their high chemo- as well as radio-resistance have been studied extensively but are still poorly understood [4,21,22]. Already in 1991, the possibility of radiotherapy promoting GBM migration and invasiveness was suggested [23]. Several studies have since then been carried out to document the effects of radiotherapy on cancer cell migration and extensive reviews of these have been done [4,5]. Majority of the results showed that ionizing radiation enhanced the migration of cancer cells including GBM [21,24–26]. A few cases [27,28] of purportedly reduced migration were included in the review [5] but it turns out that one of these [28] actually reported radiation-induced enhancement of migration and not reduction. While this apparent controversial outcome is yet to be fully resolved, the divergence of results has been putatively explained in terms of the irradiation itself (dose delivered, rate of delivery), method of assessing migration (wound healing assay, Boyden chamber), setup (*in vivo, in vitro, ex vivo*) [5] and natural heterogeneity of glioma cells [4,20]. We have therefore addressed the question whether ionizing radiation might inadvertently promote invasiveness of GBM cells via changes in cell migratory ability using experimental strategies that take the divergence of previous results into account. Regarding dose, we used 20 Gy, 10 Gy and 2 Gy to cover lethal and non-lethal regimes. Regarding measurement of migration, our method is not an end-point assessment as the Boyden chamber frequently used in previous reports, but a real time assay that measures both attachment and migration. Regarding setup and heterogeneity of glioma cells, the currently proposed need for personalized radiation medicine [4,20,21] informed our strategy, namely, an *in vitro* setup using that can be adopted for parallel monitoring of patient samples in the clinic so as to optimize treatments.

In addition to aiming at clinical translation, this work also aims at providing radiobiological phenotyping of cells based on post-irradiation and post-chemotherapy migration. We found that the highly radio-resistant cancer cell, T98G glioblastoma cell line, and the highly radio-resistant non-cancer cell, HCN2 neuronal cell line, show enhanced migration following lethal and non-lethal doses of X-rays. On the other hand, the well-known radio-responsive blood cells, precisely, HL60-derived macrophages went into cell death following irradiation. We further examined the impact of combined radiotherapy and chemotherapy (using Paclitaxel) on cell migration to guide the overall interpretation of our results. Our work sets the stage for *ex vivo* parallel monitoring of patient samples to inform personalized anti-invasion and anti-metastasis strategies for improved radiotherapy and chemotherapy outcomes.

## 2. Materials and Methods

### 2.1 Cell culture

The HCN2 cells are human brain encephalitis-derived neurons which we purchased from the American Type Culture Collection, ATCC, (HCN-2 ATCC ® CRL-10742™). We grow them in 90% DMEM with 4 mM L-glutamine adjusted to contain 1.5 g/L sodium bicarbonate, 4.5 g/L glucose, and 10% FBS, following ATCC protocols. HCN2 cells were irradiated following passages 2 to 8 and experiments stopped after passage 14. We used two cell culture incubators whereby all the cells were grown in one, while the other incubator was used for the ECIS experiments. The incubators were maintained at 95% air; 5% CO2 and a temperature of 37°C.

HL60 cells, human peripheral blood derived acute promyelocytic leukemia cancer cells, were purchased from ATCC (HL-60 ATCC ® CCL-240™) and grown in suspension using standard methods and media, namely, RPMI 1640, supplemented with 10% fetal bovine serum (FBS) and 1% Penicillin/Streptomycin. The HL60 cells were induced to differentiate into macrophages using phorbol 12-myristate 13-acetate, PMA, using the exact protocol reported in our previous works [33,34].

The T98G cells are human brain derived glioblastoma multiform cells with fibroblastic morphology [2]. We purchased T98G cells from ATCC (T98G ATCC® CRL-1690™) and grew them as prescribed using ATCC-formulated Eagle’s Minimum Essential Medium and 10% FBS. Both T98G and HCN2 cells are adherent in nature.

Cell viability was assessed using Trypan-Blue exclusion test. For irradiation and migration experiments, adherent cells were trypsinized and resuspended in culture medium at a density of about 2 to 5 x 10^5^ cells/ml in T-25 flasks. Several T-25 flasks bearing cells treated with radiotherapy and or chemotherapy were kept in the second incubator and used for viability tests and morphological imaging following irradiation/chemotherapy at 24 hours, 48 hours, 72 hours, 96 hours and 120 hours. An inverted phase-contrast fluorescent microscope (IN300-Fluor AMScope) was used for the morphological imaging.

### 2.2 Irradiation of cells

We used a dedicated benchtop compact cabinet X-ray cell irradiator, Faxitron CellRad (Tucson, AZ, USA) for delivering doses to cells. Its specifications include: Energy range, 10-130 kV; Tube current, 0.1-5 mA; Tube Power, 650 W; Dose Rate (130kVp, 5.0mA) up to 50Gy/min (unfiltered); up to: 13Gy/min (0.5mm Al filter); Focal spot size, nominal, 1.0 x 1.4 mm; Source to object distance, 17” (44 cm); Exposure time, 5 sec to 180 min (1 sec increments). We selected the autodose control of the unit and irradiated cells at a dose-rate of 0.53 Gy/min, using 100 kVp and 4 mA. Cells inside T-25 flasks were irradiated inside the X-Ray cabinet at single doses of 2, 10 and 20 Gy. Molecular level readouts for additional confirmation of irradiation effects apart from cell viability tests and morphometry included the assessment of reactive oxygen species using our recently published protocol [53].

### 2.3 Measurement of cell attachment and migration

The Electric Cell Impedance Sensing (ECIS®) device (AppliedBiophysics, New York), is a well-established real-time, label-free, impedance-based device to study the activities of cells grown in tissue culture [54,55]. These include attachment, migration, morphological changes, and other behaviors directed by the cell’s cytoskeleton. The ECIS device was used to measure cell attachment and migration which are reflected in the normalized resistance or impedance readouts. At the end of the ECIS measurement, the wells were examined to confirm the presence of cells and compare their morphology with those taken in the T-25 flasks used for morphological imaging and subsequently for morphometric analysis.

### 2.4 Morphometry and data analysis using ImageJ’s FracLac and Origin ANOVA

Phase contrast images of cells inside T-25 flasks were taken using 20x objectives at the time of irradiation/chemotherapy (0 hour), 24 hours, 48 hours, 72 hours, 96 hours, and 120 hours following irradiation/chemotherapy. The images were segmented in ImageJ (US National Institutes of Health) and its FracLac plugin was used in measuring the lacunarity and fractal dimensions of the images, to enable comparisons. Here, lacunarity quantifies gaps and heterogeneity in images. By definition, the most basic number for lacunarity, *λ*, is given as:

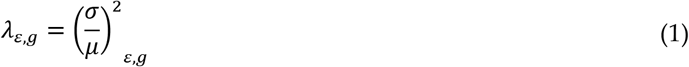

where σ is the standard deviation and μ is the mean for pixels per box at size, ε, in a box count at an orientation, g.

To make comparisons and assess statistical significance of differences in normalized resistance or impedance from ECIS measurement, viability from Trypan blue exclusion experiments and morphometry, the analysis of variance (ANOVA) function in OriginPro (OriginLab, Northampton, MA, USA) was used.

## 3. Results

### 3.1 Both radiation and chemotherapy enhance cancer cell (glioblastoma) migration before cell death

Glioblastomas, the most aggressive primary brain tumors, are highly radio-resistant and chemoresistant, yet the standard of care in the clinic is still a triple modality of surgical resection followed by radiotherapy and chemotherapy [1,29]. In spite of this triple treatment modality, median survival of patients thus treated is still limited to 14 months, even with early detection and treatment [20,30]. Such abysmal outcome confirms the aggressive and invasive nature of GBM. Instructively, 90% of GBMs relapse close (1-2 cm) to the primary tumor site following resection [31]. Both resection and targeted irradiation are believed to have limited success in eliminating tumors due to GBM diffusively invading surrounding tissues and evading and/or resisting treatment [20,32]. Even when concomitant and adjuvant chemotherapy are added to radiotherapy using drugs such as temozolomide (TMZ), local relapses occur and median survival remains 14 months compared to 12 months with RT alone [1]. Since these treatments actually remove or kill some cells, the relapses may be due to treatment-induced augmentation of migration of the GBM cells that are not immediately killed by chemo-radiotherapy. We measured the attachment and migration of T98G glioblastoma cells following radiotherapy using 20 Gy and 2 Gy to represent both lethal and non-lethal doses respectively. Notably, the dose that reduces the survival fraction of GBMs to 50%, D50, is reported to be about 2 Gy [4]. We also treated T98G cells solely with Paclitaxel (5 μM) and also concurrently with 20 Gy irradiation. The results are shown in Figure 1 as representative of all three repeats for this experiment. Figure S1 shows another of these three repeats. Cells were allowed to migrate in culture conditions for one week (160 hours). As expected, the non-irradiated cells (T98G ctl) and the cells treated with non-lethal dose of 2 Gy (T98G+2Gy) continued to migrate throughout the period of experiment (160 hours). Even the cells treated with 2 Gy showed greater diffuse migration than the untreated cells (Figure 1a). Interestingly, within the first 24 hours post-treatment, all cells treated with radiation (20 Gy and 2 Gy) and Paclitaxel (Pac), migrated more than the controls (Figure 1b). These results clearly illustrate the alteration of cancer cell migration by radiotherapy and chemotherapy prior to cell death.

**Figure 1.**
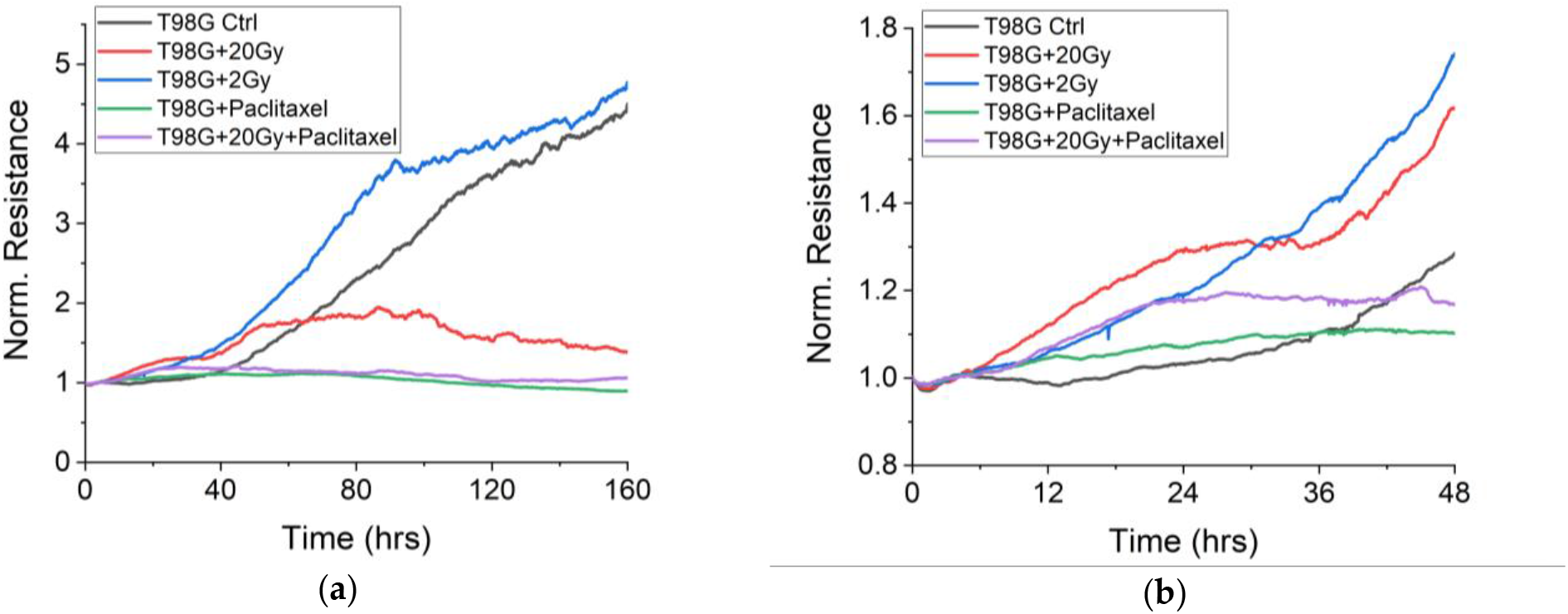
Migration of T98G cells post radiotherapy (20 Gy and 2 Gy) and chemotherapy (Paclitaxel). (a) Seven days or 160 hours of tracking. The 20 Gy treated cells eventually died just as the Paclitaxel treated cells (Pac) and the cells treated with both Pac and 20 Gy. (b) Same data as in (a) showing the first 48 hours. All irradiated cells (20 Gy and 2 Gy) migrated more than controls, with p < 0.01 (20 Gy), between 20 and 40 hours.

To corroborate the migration results, we monitored the morphology and viability of cells at 0 hour just after induction of radiotherapy and/chemotherapy, 24 hours, 48 hours, 72 hours, 96 hours and 120 hours after. Typical cell morphology 24 hours post radiotherapy and chemotherapy for experiments in Figure 1 and Figure S1 are presented in Figure S2. Typical cell morphology 96 hours post radiotherapy and chemotherapy for experiments in Figure 1 and Figure S1 are presented in Figure 2. Clearly, the much reduced attachment and reduced migration of the Paclitaxel-treated cells are reflected in the more rounded morphology and homogeneity of these cells compared to the more migratory cells, especially T98G + 2Gy and T98G controls at 96 hours and beyond (Figure 2). We have quantified these qualitative differences using lacunarity, a measure of morphological heterogeneity or gappiness in images. Typical cell morphology 48 hours after radiotherapy are shown in Figure S3 and the corresponding lacunarity results presented in Figure S4. The closer the slope of the graph of lacunarity as a function of box size is to 0, the more homogeneous the image. Obviously, cells treated with Paclitaxel only are the most homogenous, (see Figure 2d, Figure S2d and Figure S3d) and their slopes in the lacunarity plots are the closest to 0 (Figure S4 and Figure S5). Curiously, the 20 Gy plus Paclitaxel treated cells end up with the greatest heterogeneity via lacunarity measure, perhaps illustrating the more complex situation of combined radiative and chemical killing (Figure S4 and Figure S5).

**Figure 2.**
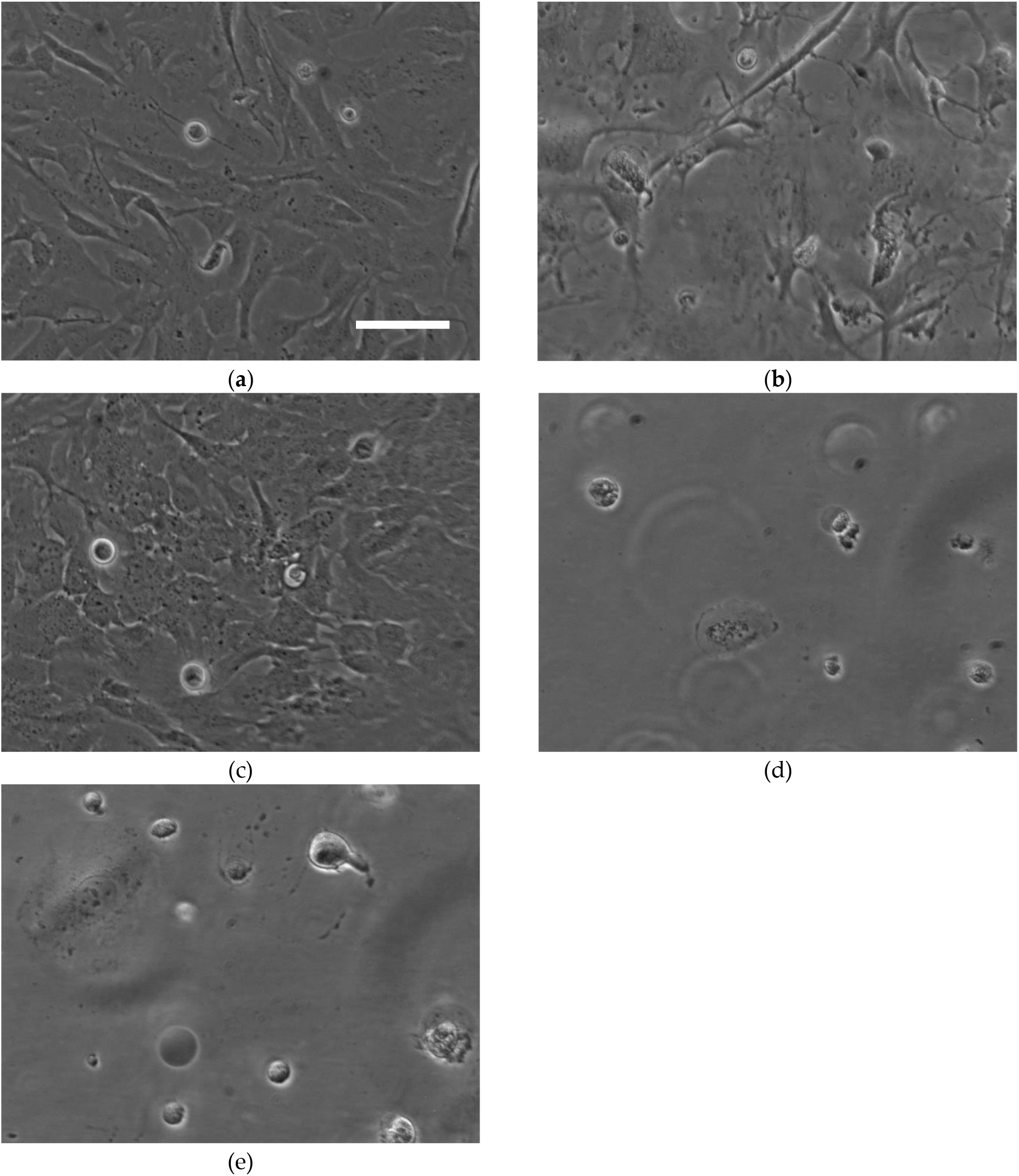
Typical cell morphology 96 hours post radiotherapy and chemotherapy for experiments in Figure 1 and Figure S1. (a) T98G Ctl. (b) T98G + 20 Gy. (c) T98G + 2 Gy. (d) T98G + Pac. (e) T98G + Pac + 20 Gy. Scale bar is 100 μm.

Furthermore, we monitored the migration of T98G cells following irradiation with 10 Gy, to have an intermediate dose for comparison with the lethal and non-lethal, and for comparison with previously published work. The dose that reduces the survival fraction to 10% or D10 is reported to be between 5 and 8 Gy [4]. Figure 3 shows that within the first 24 hours following irradiation with 10 Gy, there is also enhancement of migration of T98G cells as with 20 Gy and 2 Gy.

**Figure 3.**
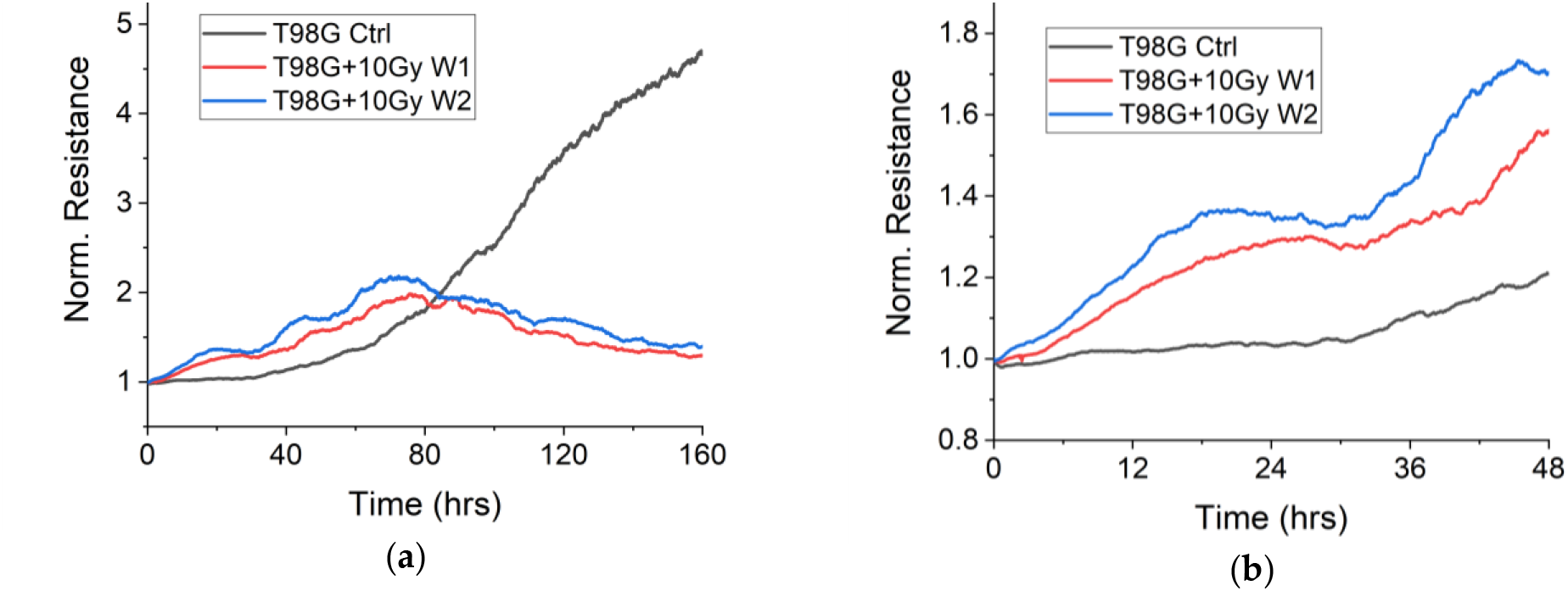
Migration of T98G cells post radiotherapy (10 Gy). (a) Seven days or 160 hours of tracking. The 10 Gy treated cells do not die within 160 hours but the irradiated cells in wells 1 and 2 do. (b) Same data as in (a) showing the first 48 hours. All irradiated cells (10 Gy) migrated more than controls, with p < 0.05 between 20 and 40 hours. The reproducibility of results is remarkable.

Remarkably, the two wells of 10 Gy irradiated cells (W1 and W2) of Figure 3 are very reproducible. To further generalize the features of Figure 3, we present in Figure 4, a fourth repeat of the experiments of Figure 3. All experiments with 10 Gy (Figure 3 and Figure 4) show two stages of enhanced migration in the first 48 hours post-irradiation, namely, a first period of very rapid migration up to 24 hours, a slowing down and another rapid regime between 30 to 48 hours. These features could be important at the molecular level, in view of developing anti-metastasis strategies for improved GBM treatments. Hence, we carried out further experiments with 2 Gy using multiple wells to focus on the migratory timescales and features as in the case of 10 Gy and 20 Gy.

**Figure 4.**
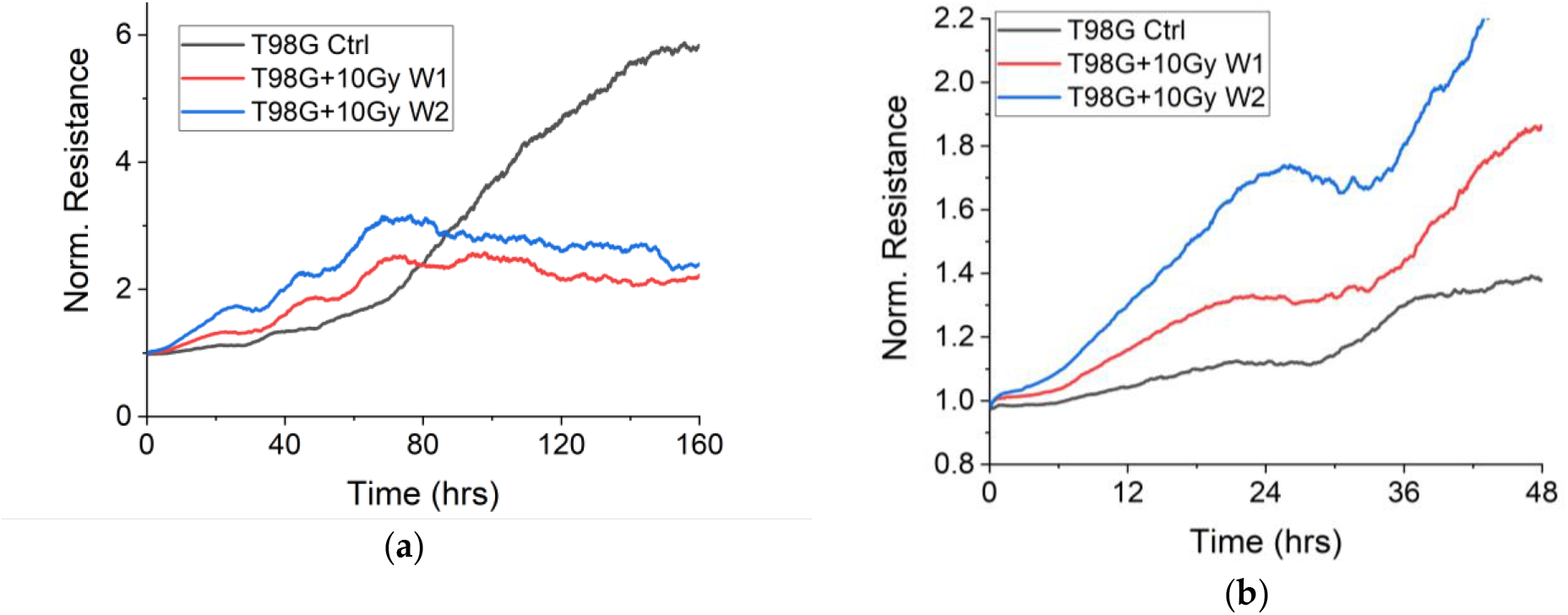
Migration of T98G cells post radiotherapy (10 Gy). (a) Seven days or 160 hours of tracking. The untreated cells do not die within 160 hours but the irradiated cells in wells 1 and 2 do. (b) Same data as in (a) showing the first 48 hours. All irradiated cells (10 Gy) migrated more than controls. This is a 4th repeat experiment enabling generalization of post-irradiation migration patterns.

Figure 5 shows that the two distinct phases of enhanced migration found with 10 Gy irradiation (and with 20 Gy irradiation), are absent for the 2 Gy irradiation in all wells of the two different experiments shown. In typical clinical irradiation, tumor cells would get varying doses in spite of uniform dose delivery to the planned tumor volume, owing to inherent heterogeneity of the tumor volume and the cells themselves.

**Figure 5.**
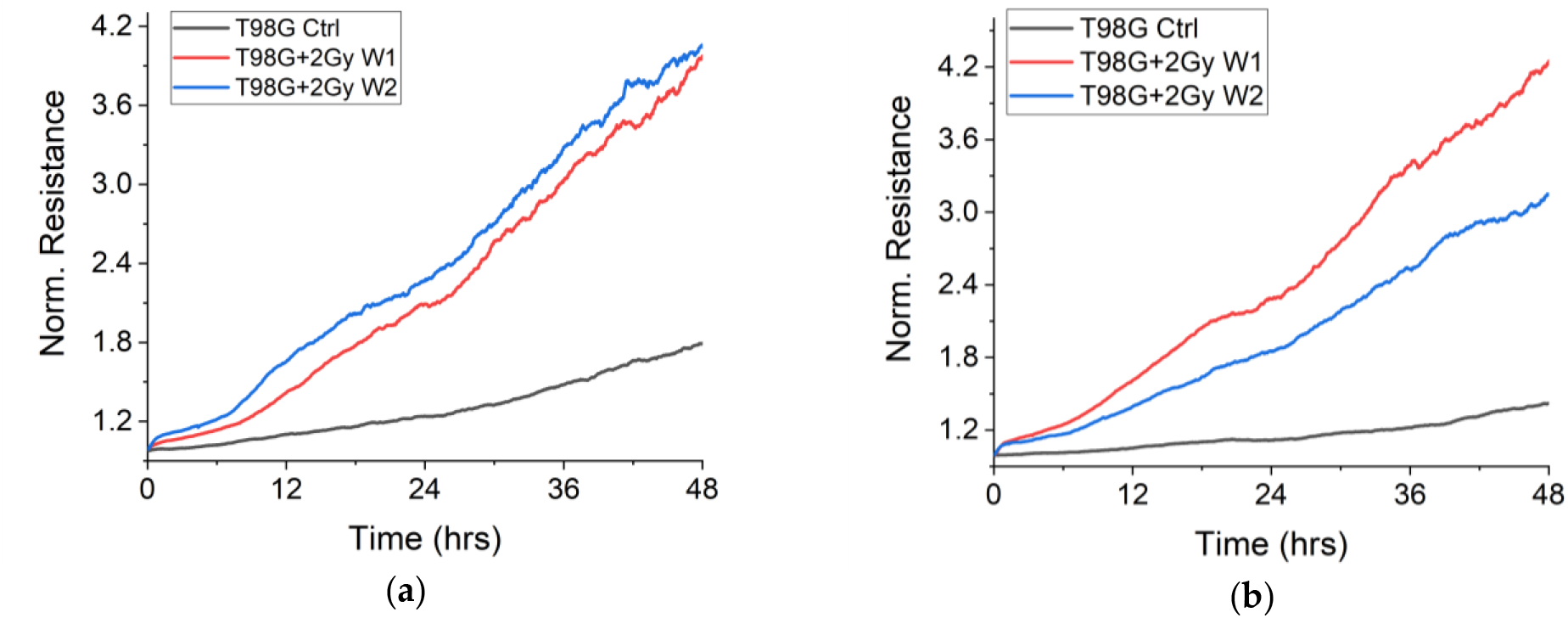
Migration of T98G cells post radiotherapy (2 Gy). (a) First 48 hours for experiment N = 4. (b) First 48 hours for experiment N = 5. The distinct phases of enhanced migration found in 10 Gy are absent with 2 Gy.

Furthermore, some non-cancer cells do get irradiated even though irradiation is usually optimized to target cancer cells and spare healthy tissue. Hence, we considered non-cancer cells of the central nervous system, CNS, that might also receive radiation in the course of GBM treatment.

### 3.2 Irradiated non-cancer cells (HCN2 neurons) attach and migrate more than un-irradiated neurons

We find that the 20 Gy irradiated HCN2 cells attach and migrate more than non-irradiated cells in the first 20 hours post irradiation (Figure 6).

**Figure 6.**
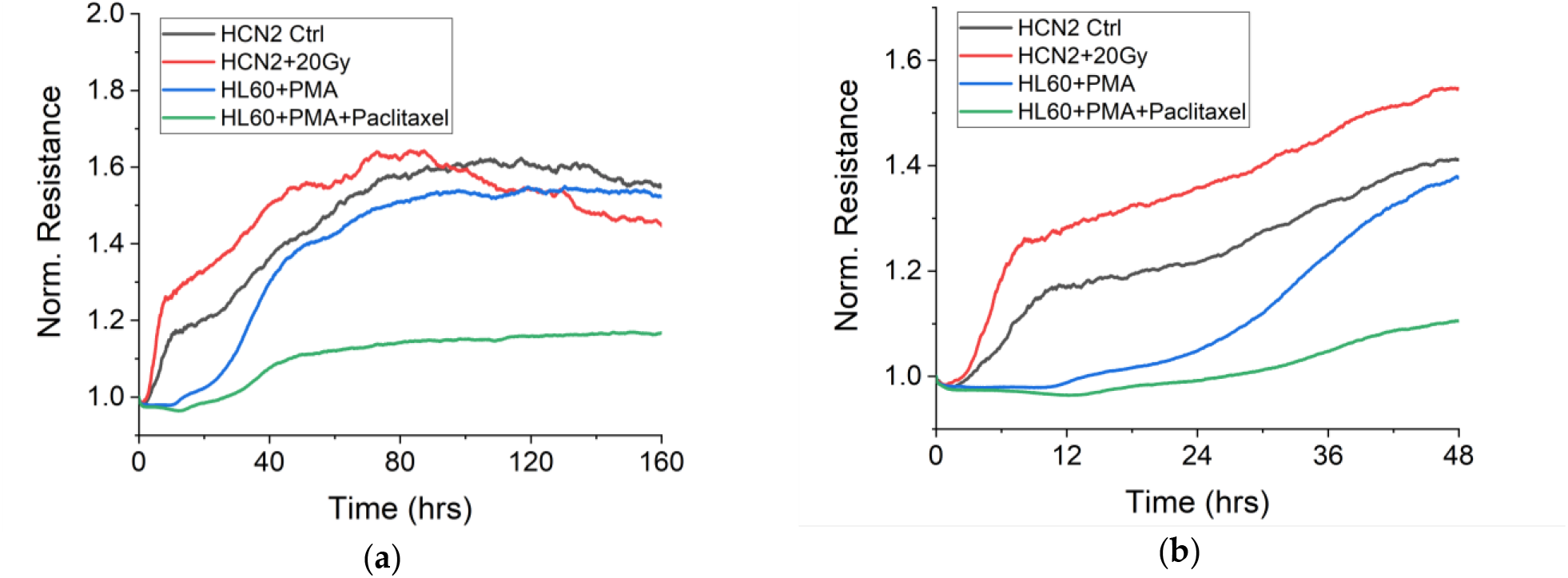
Migration of HCN2 neurons post-radiotherapy (20 Gy) and HL60-derived macrophages post-chemotherapy (Paclitaxel). (a) Seven days or 160 hours of tracking. The 20 Gy treated HCN2 cells eventually go into senescence while Pac-treated macrophages responded to treatment by migrating less than untreated macrophages throughout. (b). The first 48 hours of data shown in (a). Irradiated HCN2 cells migrated more than unirradiated HCN2 cells. HL60 derived macrophages treated with Paclitaxel migrated less than untreated cells.

The enhanced migration of non-cancer brain cells (here, neurons) in the context of radiotherapy implies that at the molecular level, the radiation-induced alteration of cell migration is much more general and fundamental. It is not merely a cancer-specific phenomenon. This implies that any non-cancer tissue that is part of the planned tumor volume, or any healthy tissue that gets irradiated in the course of treatment, may also be altered with respect to the cytoskeleton and migration. Additionally, to ascertain the reliability of the ECIS tracking of migration via impedance or resistance, we induced HL60 cells to differentiate into macrophages using well established protocols [33,34]. Glial cells or microglia in the brain are resident macrophages and macrophages are among the most migratory cells of the body [8]. Our ECIS system showed that the HL60-derived macrophages migrated in a pattern very similar to the HCN2 neuronal cells (Figure 6). The Paclitaxel treated macrophages migrated less than the untreated cells, throughout the course of the experiment (Figure 6), in contrast to the highly radio-resistant and chemo-resistant GBM cells (T98G) which migrated more in the first 24 hours following Paclitaxel treatment. This migration result reflects the known high chemo-resistance of T98G cells.

## 4. Discussion

### 4.1 Alteration of cell migration, cancer metastasis and physics of cancer

Our working hypothesis that radiotherapy and chemotherapy may alter migration of both cancer and non-cancer cells has been confirmed by the results presented in this work, with some qualifications. The overall alteration is an enhancement of migration, with statistical significance of the enhancement increasing with radiation dose in the case of radiotherapy. How this alteration of migration might affect metastasis and cancer relapse is the main clinical implication that needs to be demonstrated. We posit that an important link between cell migration and cancer metastasis, that needs to be explored, is the cytoskeleton of cells. We again note that migration is a crucial step in the metastatic cascade [15,16]. To migrate, cells actively alter their mechanical properties through their actin-myosin cytoskeleton [35]. We have recently shown *in vitro* [36] and others have shown in patient samples [11,13,37] that chemotherapy inadvertently alters the mechanical properties of leukemic cancer cells in pro-metastatic ways. Our present results showing that chemotherapy (here Paclitaxel) alters the migration of brain cancer cells and neurons suggest that this alteration happens via changes to the actin cytoskeleton of cells.

Moreover, the alteration of migration of both cancer cells and non-cancer cells by radiotherapy, which we have demonstrated here, suggests that the cytoskeletal properties of cells are being altered by the X-ray photon interactions. Interestingly a recent report showed that non-lethal single dose radiation (2 Gy) influenced the invasiveness, cell stiffness and actin cytoskeleton properties of two GBM cell lines, namely, LN229 and U87 [24]. Although the authors reported reduced invasiveness following the 2 Gy irradiation, they found “generalized stiffness” to be a profound marker of the invasiveness of a tumor cell population [24]. This finding by itself is not new. Cellular mechanical properties have been shown to be good markers for metastatic potential of cancer cells [38,39] and can be used for diagnosis of cancer itself [15,40,41]. Furthermore, chemotherapeutic drugs alter the mechanical properties of cells during cancer treatment [37]. Following recent realization that key stages in the metastatic cascade such as migration, involve the mechanical and physical properties of cells [42], physical oncology or physics of cancer [16,42,43] has emerged, clearly illustrating the connection between cell mechanics, cell migration and metastasis.

Our present results occasion the need to extensively address the impact of radiotherapy on the physics of cancer, that is, the connection between radiotherapy, cell migration and cell mechanical properties, in the context of metastasis. Other results that suggest the need to consider the impact of radiotherapy on cell mechanics and migration include the report that low doses of ionizing radiation (0.8 Gy or less) can promote tumor growth and metastasis by enhancing endothelial cell migration and angiogenesis [6], the investigation which focused on the effects of ionizing radiation on the cytoskeleton of endothelial cells and endothelial monolayer permeability [44], those that measured impact of X-rays and heavy ions on neural cell mechanics [45], and the measurement of mechanical properties of human umbilical vein endothelial cells following irradiation [46]. The further experiments that we suggest here (impact of radiotherapy on cell mechanics in view of cancer metastasis) will benefit from the recent comparison and notes for selection of techniques for cell mechanics in view of clinical applications [47] and the emergence of very rapid techniques for mechanical phenotyping [48]. An important aspect of our results in this work which call for careful selection of techniques for measurement and intervention is the time scale of events being measured. We discuss time scales next.

### 4.2 Timescales in alteration of migration, reversible intervention and limitations

The proliferation, attachment and migration of cells are all phenomena that happen in hours. Hence, different cell types have different doubling times. Our results based on attachment and migration of various cells types (T98G Glioblastoma, HCN2 neurons, HL60-derived macrophages) should indeed show different baseline readouts of migration for the control cells of each lineage, and this indeed is the case. However, with radiotherapy and chemotherapy, the time-scales of radiation-induced and/or drug-induced effects should become apparent. Since our migration assay provides continuous readouts throughout the week-long experiments, we have found time-scales involved in the induced effects. With radiotherapy at 10 Gy and 20 Gy, there are two distinct phases in the migration: an initial high rate of migration for about 24 hours following treatment. Our results suggest a novel timescale that can be targeted by parallel anti-migratory drugs, especially those that have reversible action such as certain cytochalasins [49,50], during radiotherapy and chemotherapy. Interestingly, anti-migratory drugs have recently been suggested by others for the improvement of radiotherapy [17,25] and chemotherapy [51,52]. Based on our results, the first 24 to 48 hours should be prime time to use the anti-migratory strategies. Obviously, reversal of the anti-migratory effects becomes necessary since the inhibition of migration by alteration of the actin cytoskeleton might induce collateral impairment of immune functions [35].

Beyond the timescales based on our 10 Gy and 20 Gy results, the 2 Gy-induced enhancement of migration fits well with the overall outlook that cells surviving radiation demonstrate increased migration/invasion and therapeutic resistance [26]. Previous reports indicating reduced migration may have been partially due to end-point assays missing the timescales of the enhanced migration that we have found in this work. The question remains whether our *in vitro* results reflect *in vivo* situation. Another limitation of our work is the 2D state of our migration assay. Migration of cells in the body is a 3D phenomenon and closer mimicking of this *in vivo* 3D situation is needed as future work. Finally, there is the limitation imposed by GBM immortalized cell lines lacking some of the properties of GBM cells found in patients [20]. However, the set-up we have used allows for patient-specific samples to be examined in order to determine new or further course of treatment, along the lines of personalized radiation medicine [4,22]. Whether radiotherapy and/or chemotherapy enhance(s) cancer cell migration in patients or not, strategies that reduce migration are already promising anti-metastasis treatments [21,26,27] which could in turn improve outcomes for patients. Our work gives timescales when such strategies might be most effective.

### 5. Conclusions

We have addressed a longstanding question whether radiotherapy might promote metastasis by enhancing one of its crucial steps, cell migration. Our results show that doses of X-rays within the range used in radiotherapy and chemotherapy alter the migration of both cancer cells (brain cancer) and non-cancer cells (neurons) prior to cell death. The alteration is an enhancement of migration, which increases in statistical significance in a dose-dependent manner. These results suggest that cell migration should be a therapeutic target in anti-metastasis strategies for improved radiotherapy and chemotherapy outcomes. Moreover, our *in vitro* setup for tracking migration presents itself as a tool that can be translated to the clinic to aid in the development of personalized radiation medicine. The heterogeneity of gliomas and the uniqueness of individual medical histories and prognosis call for such personalization. For a robust translational tool, 3D tissue constructs and scaffolds could be added to the 2D migration and morphometric assay presented here. This work presents *in vitro* evidence for possible *in vivo* alterations of cell migration by chemotherapy and radiotherapy, leading to poor outcomes for patients. It provides a scientific basis for formulating anti-metastasis drugs or strategies that reversibly reduce migration within 48 hours of treatment. It also engenders the feasibility of migration-based *ex vivo* monitoring of patient-specific cells to customize and optimize radiotherapy and chemotherapy for improved outcomes especially in cases of brain cancers.

## Supplementary Materials

The following are in the Supplements.

Figure S1. Repeat Experiment for Migration of T98G cells post radiotherapy (20 Gy and 2 Gy) and chemotherapy (Paclitaxel).

Figure S2. Typical cell morphology 24 hours post radiotherapy and chemotherapy for experiments in Figure 1 and Figure S1.

Figure S3. Typical cell morphology 48 hours post radiotherapy and chemotherapy for experiments in Figure 1 and Figure S1.

Figure S4. Typical lacunarity-based cell morphometry 48 hours post radiotherapy and chemotherapy for experiments in Figure 1 and Figure S1.

Figure S5. Typical lacunarity-based cell morphometry 96 hours post radiotherapy and chemotherapy for experiments in Figure 1 and Figure S1.

## Author Contributions

**C**onceptualization, supervision, project administration, funding acquisition, and writing—original draft preparation, A.E.; Methodology, data analysis, A.E., M.M., A.B., M.J.M; Investigation, experiments, M.M., M.J.M, C.W., H.A., A.B., A.W., C.N., J.B., H.D., K.A., AE; Writing—review and editing, A.E., M.M., M.J.M, H.D.

## Acknowledgments

The authors are grateful to Dr Andrew Baruth and Dr Michael Nichols for allowing us the use of some laboratory equipment and space, especially additional biosafety cabinets and incubators.

## References

[1] J. Mann, R. Ramakrishna, R. Magge, A.G. Wernicke, Advances in radiotherapy for glioblastoma, Front Neurol. 8 (2018) 1–11. doi:10.3389/fneur.2017.00748.

[2] H. Murad, Y. Alghamian, A. Aljapawe, A. Madania, Effects of ionizing radiation on the viability and proliferative behavior of the human glioblastoma T98G cell line, BMC Res Notes. 11 (2018) 1–6. doi:10.1186/s13104-018-3438-y.

[3] H.E. Barker, J.T.E. Paget, A.A. Khan, K.J. Harrington, The Tumour Microenvironment after Radiotherapy: Mechanisms of Resistance and Recurrence, Nat Rev Cancer. 15 (2016) 409–425. doi:10.1038/nrc3958.The.

[4] C. Zimmer, S. Bette, M. Wank, M. Barz, B. Wiestler, K. Kessel, F. Liesche, B. Meyer, T. Schmid, S. Combs, J. Schlegel, J. Gempt, D. Schilling, Human Glioma Migration and Infiltration Properties as a Target for Personalized Radiation Medicine, Cancers (Basel). 10 (2018) 456. doi:10.3390/cancers10110456.

[5] C. Moncharmont, A. Levy, J.B. Guy, A.T. Falk, M. Guilbert, J.C. Trone, G. Alphonse, M. Gilormini, D. Ardail, R.A. Toillon, C. Rodriguez-Lafrasse, N. Magné, Radiation-enhanced cell migration/invasion process: A review, Crit Rev Oncol Hematol. 92 (2014) 133–142. doi:10.1016/j.critrevonc.2014.05.006.

[6] I. Sofia Vala, L.R. Martins, N. Imaizumi, R.J. Nunes, J. Rino, F. Kuonen, L.M. Carvalho, C. Rüegg, I.M. Grillo, J.T. Barata, M. Mareel, S.C.R. Santos, Low doses of ionizing radiation promote tumor growth and metastasis by enhancing angiogenesis, PLoS One. 5 (2010). doi:10.1371/journal.pone.0011222.

[7] P. Mehlen, A. Puisieux, Metastasis: a question of life or death, Nat Rev Cancer. 6 (2006) 449–458. doi:10.1038/nrc1886.

[8] T.N. Seyfried, L.C. Huysentruyt, On the origin of cancer metastasis., Crit Rev Oncog. 18 (2013) 43–73.

[9] P.S. Steeg, Tumor metastasis: mechanistic insights and clinical challenges, Nat Med. 12 (2006) 895–904. doi:10.1038/nm1469.

[10] Y.A. Fouad, C. Aanei, Revisiting the hallmarks of cancer., Am J Cancer Res. 7 (2017) 1016–1036.

[11] S. Ran, The role of TLR4 in chemotherapy-driven metastasis, Cancer Res. 75 (2015) 2405–2410. doi:10.1158/0008-5472.CAN-14-3525.

[12] G.F. Weber, Why does cancer therapy lack effective anti-metastasis drugs?, Cancer Lett. 328 (2013) 207–11. doi:10.1016/j.canlet.2012.09.025.

[13] L. Volk-Draper, K. Hall, C. Griggs, S. Rajput, P. Kohio, D. DeNardo, S. Ran, Paclitaxel therapy promotes breast cancer metastasis in a TLR4-dependent manner., Cancer Res. 74 (2014) 5421–34. doi:10.1158/0008-5472.CAN-14-0067.

[14] A.A. Geldof, B.R. Rao, Doxorubicin treatment increases metastasis of prostate tumor (R3327-MatLyLu)., Anticancer Res. 8 (1988) 1335–9.

[15] S. Suresh, Biomechanics and biophysics of cancer cells., Acta Biomater. 3 (2007) 413–438.

[16] N.M. Moore, L.A. Nagahara, Physical biology in cancer. 1. Cellular physics of cancer metastasis., Am J Physiol Cell Physiol. 306 (2014) C78–9. doi:10.1152/ajpcell.00292.2013.

[17] M.A. Grotzer, A. Neve, M. Baumgartner, Dissecting brain tumor growth and metastasis in vitro and ex vivo, J Cancer Metastasis Treat. 2 (2016) 149. doi:10.20517/2394-4722.2016.02.

[18] D. Hanahan, R.A. Weinberg, The hallmarks of cancer., Cell. 100 (2000) 57–70.

[19] Y.-N. Liu, B.-B. Kang, J.H. Chen, Transcriptional regulation of human osteopontin promoter by C/EBPalpha and AML-1 in metastatic cancer cells., Oncogene. 23 (2004) 278–88. doi:10.1038/sj.onc.1207022.

[20] S. Caragher, A.J. Chalmers, N. Gomez-Roman, Glioblastoma’s Next Top Model: Novel Culture Systems for Brain Cancer Radiotherapy Research, Cancers (Basel). 11 (2019) 44. doi:10.3390/cancers11010044.

[21] J.P. Zepecki, K.M. Snyder, M.M. Moreno, E. Fajardo, A. Fiser, J. Ness, A. Sarkar, S.A. Toms, N. Tapinos, Regulation of human glioma cell migration, tumor growth, and stemness gene expression using a Lck targeted inhibitor, Oncogene. (2018). doi:10.1038/s41388-018-0546-z.

[22] I. Manini, F. Caponnetto, A. Bartolini, T. Ius, L. Mariuzzi, C. Di Loreto, A.P. Beltrami, D. Cesselli, Role of microenvironment in glioma invasion: What we learned from in vitro models, Int J Mol Sci. 19 (2018). doi:10.3390/ijms19010147.

[23] C. von Essen, Radiation enhancement of metastasis: a review., Clin Exp Metastasis. 9 (1991) 77–104.

[24] D. Vordermark, C. Vogel, F. Dehghani, M. Bache, T. Hohmann, S. Ensminger, C. Ghadban, U. Grabiec, The Impact of Non-Lethal Single-Dose Radiation on Tumor Invasion and Cytoskeletal Properties, Int J Mol Sci. 18 (2017) 2001. doi:10.3390/ijms18092001.

[25] N. Becker, F.L.R. Ferreira, M. Flentje, D. Sisario, V. Fiedler, H. Zimmermann, V.L. Sukhorukov, C.S. Djuzenova, A. Katzer, S. Memmel, C. Zöller, R. Heiden, L. Eing, M. Sauer, Migration pattern, actin cytoskeleton organization and response to PI3K-, mTOR-, and Hsp90-inhibition of glioblastoma cells with different invasive capacities, Oncotarget. 8 (2017) 45298–45310. doi:10.18632/oncotarget.16847.

[26] F.B. Furnari, A. Purves, S.K. De, B. Hu, B. Wu, D. Sarkar, L. Emdad, S.K. Das, M. Pellecchia, P.B. Fisher, K. Valerie, T.P. Kegelman, W.K. Cavenee, S. Talukdar, J.M. Beckta, J. Wei, Inhibition of radiation-induced glioblastoma invasion by genetic and pharmacological targeting of MDA-9/Syntenin, Proc Natl Acad Sci. 114 (2016) 370–375. doi:10.1073/pnas.1616100114.

[27] F. Fehlauer, M. Muench, E. Richter, D. Rades, The inhibition of proliferation and migration of glioma spheroids exposed to temozolomide is less than additive if combined with irradiation., Oncol Rep. 17 (2007) 941–5.

[28] V.R. Gogineni, A.K. Nalla, R. Gupta, M. Gujrati, J.D. Klopfenstein, S. Mohanam, J.S. Rao, A3B1 Integrin Promotes Radiation-Induced Migration of Meningioma Cells, Int J Oncol. 38 (2011) 1615–1624. doi:10.3892/ijo.2011.987.

[29] A.A. Thomas, M.S. Ernstoff, C.E. Fadul, Immunotherapy for the treatment of glioblastoma, Cancer J. 18 (2012) 59–68. doi:10.1097/PPO.0b013e3182431a73.

[30] I. Paw, R.C. Carpenter, K. Watabe, W. Debinski, H.W. Lo, Mechanisms regulating glioma invasion, Cancer Lett. 362 (2015) 1–7. doi:10.1016/j.canlet.2015.03.015.

[31] C. Hess, J. Schaaf, R. Kortmann, M. Schabet, M. Bamberg, Malignant glioma: patterns of failure following individually tailored limited volume irradiation., Radiother Oncol. 30 (1994) 146–9.

[32] E.C. Holland, Glioblastoma multiforme: The terminator, Proc Natl Acad Sci. 97 (2002) 6242–6244. doi:10.1073/pnas.97.12.6242.

[33] A.E. Ekpenyong, U.F. Keyser, J. Guck, C. Fiddler, E.R. Chilvers, G. Whyte, S. Paschke, K. Chalut, S. Pagliara, F. Lautenschläger, Viscoelastic Properties of Differentiating Blood Cells Are Fate- and Function-Dependent, PLoS One. 7 (2012) e45237. doi:10.1371/journal.pone.0045237.

[34] C.J. Chan, A.E. Ekpenyong, S. Golfier, W. Li, K.J. Chalut, O. Otto, J. Elgeti, J. Guck, F. Lautenschl?ger, Myosin II activity softens cells in suspension, Biophys J. 108 (2015). doi:10.1016/j.bpj.2015.03.009.

[35] S.M. Man, A. Ekpenyong, P. Tourlomousis, S. Achouri, E. Cammarota, K. Hughes, A. Rizzo, G. N. Ng, J.A. Wright, P. Cicuta, J.R. Guck, C.E. Bryant, Actin polymerization as a key innate immune effector mechanism to control Salmonella infection, Proc Natl Acad Sci U S A. 111 (2014). doi:10.1073/pnas.1419925111.

[36] S. V. Prathivadhi-Bhayankaram, J. Ning, M. Mimlitz, C. Taylor, E. Gross, M. Nichols, J. Guck, A.E. Ekpenyong, Chemotherapy impedes in vitro microcirculation and promotes migration of leukemic cells with impact on metastasis, Biochem Biophys Res Commun. 479 (2016) 841–846. doi:10.1016/j.bbrc.2016.09.121.

[37] W.A. Lam, M.J. Rosenbluth, D.A. Fletcher, Chemotherapy exposure increases leukemia cell stiffness., Blood. 109 (2007) 3505–8. doi:10.1182/blood-2006-08-043570.

[38] V. Swaminathan, K. Mythreye, E.T. O’Brien, A. Berchuck, G.C. Blobe, R. Superfine, Mechanical stiffness grades metastatic potential in patient tumor cells and in cancer cell lines., Cancer Res. 71 (2011) 5075–80. doi:10.1158/0008-5472.CAN-11-0247.

[39] J. Guck, S. Schinkinger, B. Lincoln, F. Wottawah, S. Ebert, M. Romeyke, D. Lenz, H.M. Erickson, R. Ananthakrishnan, D. Mitchell, J. Käs, S. Ulvick, C. Bilby, Optical deformability as an inherent cell marker for testing malignant transformation and metastatic competence, Biophys J. 88 (2005) 3689–98. doi:10.1529/biophysj.104.045476.

[40] T.W. Remmerbach, F. Wottawah, J. Dietrich, B. Lincoln, C. Wittekind, J. Guck, Oral cancer diagnosis by mechanical phenotyping, Cancer Res. 69 (2009) 1728–32. doi:10.1158/0008-5472.CAN-08-4073.

[41] H.T.K. Tse, D.R. Gossett, Y.S. Moon, M. Masaeli, M. Sohsman, Y. Ying, K. Mislick, R.P. Adams, J. Rao, D. Di Carlo, Quantitative diagnosis of malignant pleural effusions by single-cell mechanophenotyping., Sci Transl Med. 5 (2013) 212ra163. doi:10.1126/scitranslmed.3006559.

[42] D. Wirtz, K. Konstantopoulos, P.C. Searson, The physics of cancer: the role of physical interactions and mechanical forces in metastasis, Nat Rev Cancer. 11 (2011) 512–522. doi:10.1038/nrc3080.

[43] R. Munbodh, A. Jackson, Quantifying cell migration distance as a contributing factor to the development of rectal toxicity after prostate radiotherapy, Med Phys. 41 (2014) 021724. doi:10.1118/1.4852955.

[44] D. Gabryś, O. Greco, G. Patel, K.M. Prise, G.M. Tozer, C. Kanthou, Radiation effects on the cytoskeleton of endothelial cells and endothelial monolayer permeability., Int J Radiat Oncol Biol Phys. 69 (2007) 1553–62. doi:10.1016/j.ijrobp.2007.08.039.

[45] Y.T. Du, J. Zhang, Q. Zheng, M.X. Li, Y. Liu, B.P. Zhang, B. Liu, H. Zhang, G.Y. Miao, Heavy ion and X-ray irradiation alter the cytoskeleton and cytomechanics of cortical neurons, Neural Regen Res. 9 (2014) 1129–1137. doi:10.4103/1673-5374.135315.

[46] A. Mohammadkarim, M. Tabatabaei, A. Parandakh, M. Mokhtari-Dizaji, M. Tafazzoli-Shadpour, M.-M. Khani, Radiation therapy affects the mechanical behavior of human umbilical vein endothelial cells, J Mech Behav Biomed Mater. 85 (2018) 188–193. doi:10.1016/J.JMBBM.2018.06.009.

[47] P.-H. Wu, D.R.-B. Aroush, A. Asnacios, W.-C. Chen, M.E. Dokukin, B.L. Doss, P. Durand-Smet, A. Ekpenyong, J. Guck, N.V. Guz, P.A. Janmey, J.S.H. Lee, N.M. Moore, A. Ott, Y.-C. Poh, R. Ros, M. Sander, I. Sokolov, J.R. Staunton, N. Wang, G. Whyte, D. Wirtz, A comparison of methods to assess cell mechanical properties, Nat Methods. (2018). doi:10.1038/s41592-018-0015-1.

[48] O. Otto, P. Rosendahl, A. Mietke, S. Golfier, C. Herold, D. Klaue, S. Girardo, S. Pagliara, A. Ekpenyong, A. Jacobi, M. Wobus, N. T?pfner, U.F. Keyser, J. Mansfeld, E. Fischer-Friedrich, J. Guck, Real-time deformability cytometry: On-the-fly cell mechanical phenotyping, Nat Methods. 12 (2015). doi:10.1038/nmeth.3281.

[49] I. Foissner, G.O. Wasteneys, Wide-ranging effects of eight cytochalasins and latrunculin A and B on intracellular motility and actin filament reorganization in characean internodal cells, Plant Cell Physiol. 48 (2007) 585–597. doi:10.1093/pcp/pcm030.

[50] M. Trendowski, Using cytochalasins to improve current chemotherapeutic approaches., Anticancer Agents Med Chem. 15 (2015) 327–35. doi:10.2174/1871520614666141016164335.

[51] S. Kumari, A.K. Badana, G.M. Mohan, G. Shailender Naik, R.R. Malla, Synergistic effects of coralyne and paclitaxel on cell migration and proliferation of breast cancer cells lines, Biomed Pharmacother. 91 (2017) 436–445. doi:10.1016/j.biopha.2017.04.027.

[52] C. Decaestecker, O. Debeir, P. Van Ham, R. Kiss, Can anti-migratory drugs be screened in vitro? A review of 2D and 3D assays for the quantitative analysis of cell migration, Med Res Rev. 27 (2007) 149–176. doi:10.1002/med.20078.

[53] B.H. Lee, S. Suresh, A. Ekpenyong, Fluorescence intensity modulation of CdSe/ZnS quantum dots assesses ROS during chemotherapy and radiotherapy for cancer cells, J Biophotonics. (2018) e201800172. doi:10.1002/jbio.201800172.

[54] I. Giaever, C.R. Keese, Micromotion of mammalian cells measured electrically., Proc Natl Acad Sci U S A. 88 (1991) 7896–900. doi:10.1073/PNAS.88.17.7896.

[55] R. Szulcek, H.J. Bogaard, G.P. van Nieuw Amerongen, Electric Cell-substrate Impedance Sensing for the Quantification of Endothelial Proliferation, Barrier Function, and Motility, J Vis Exp. (2014) 1–12. doi:10.3791/51300.

