## Supplementary Materials for "Radiotherapy and chemotherapy alter migration of brain cancer cells before cell death"

| 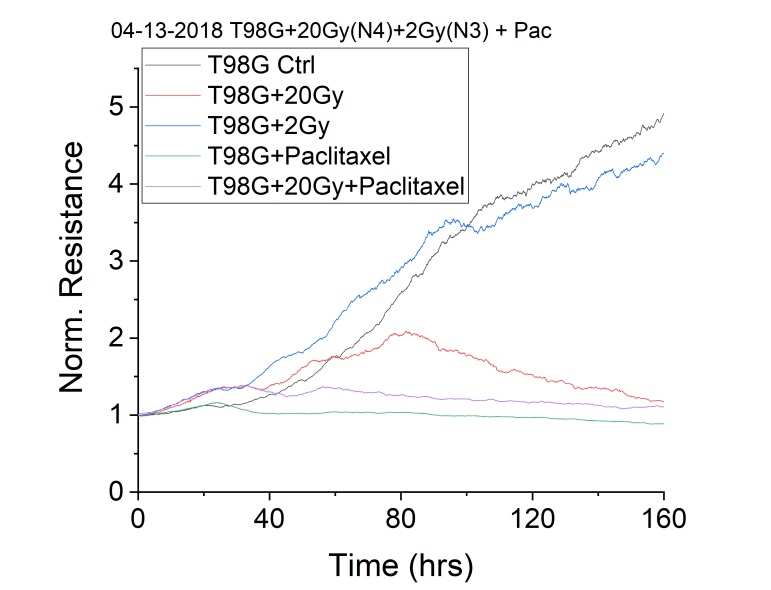  (**a**) | 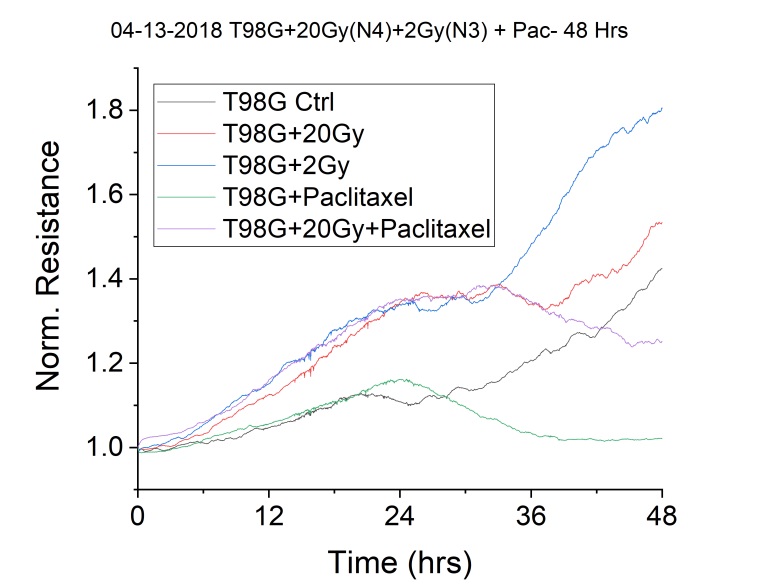  (**b**) |
| --- | --- |

**Figure S1.** Repeat experiment for migration of T98G cells post radiotherapy (20 Gy and 2 Gy) and chemotherapy (Paclitaxel). (a) Seven days or 160 hours of tracking. The 20 Gy treated cells eventually died just as the Paclitaxel treated cells (Pac) and the cells treated with both Pac and 20 Gy. (b) Same data as in (a) showing the first 48 hours. All irradiated cells (20 Gy and 2 Gy) as well as Pac-treated cells migrated more than controls. N1 is shown in Figure 1 and N2 is shown here. N3 is similar.

| 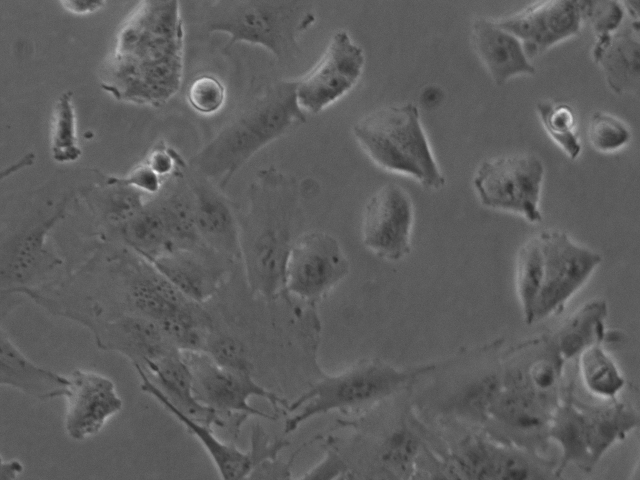  (**a**)  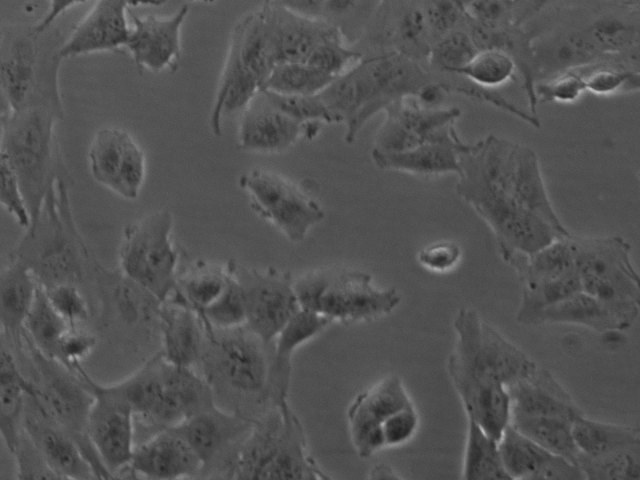  (c)  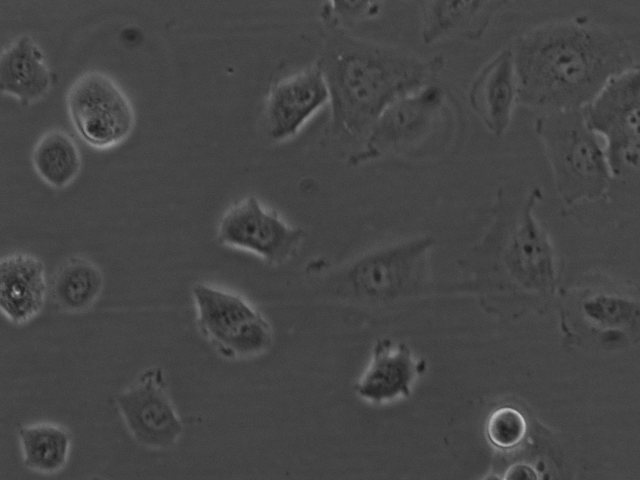  (e) | 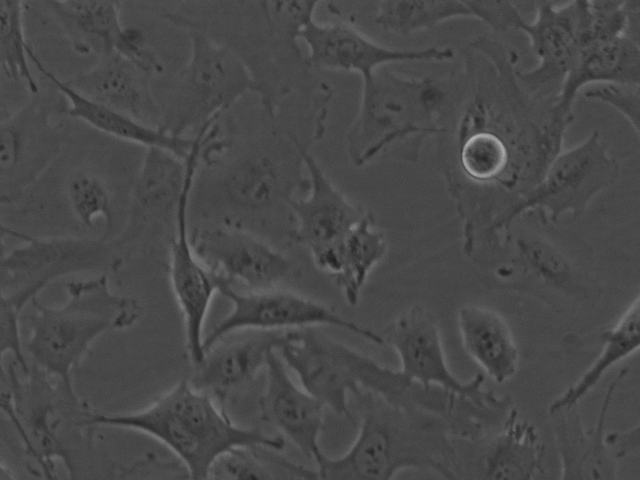  (**b**)  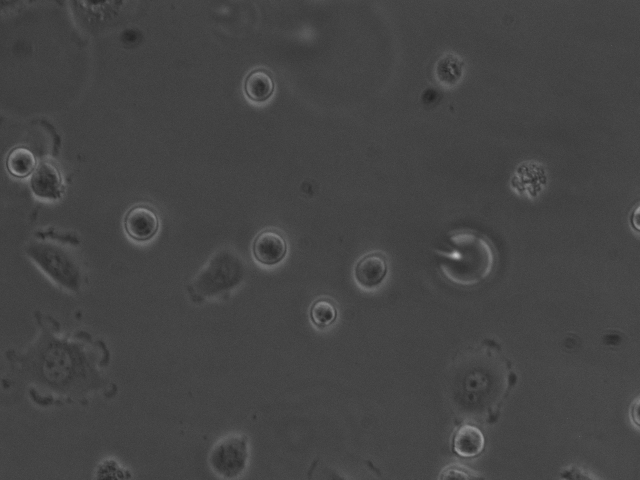  (d) |
| --- | --- |

**Figure S2.** Typical cell morphology 24 hours post radiotherapy and chemotherapy for experiments in Figure 1 and Figure S1. (a) T98G Ctl. (b) T98G + 20 Gy. (c) T98G + 2 Gy. (d) T98G + Pac. (e) T98G + Pac + 20 Gy. Scale bar is 100 μm.

| 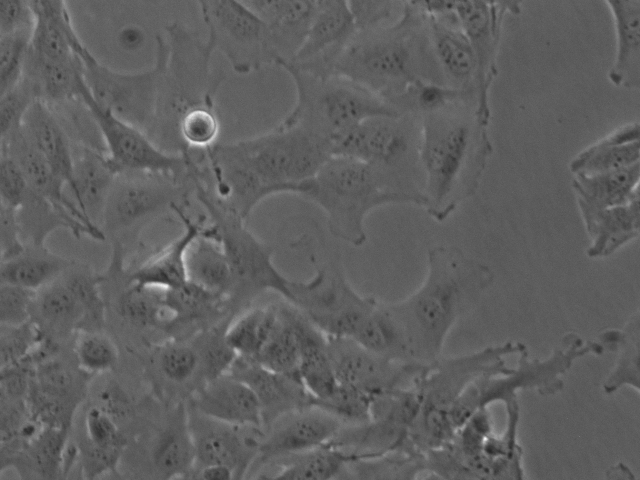  (**a**)  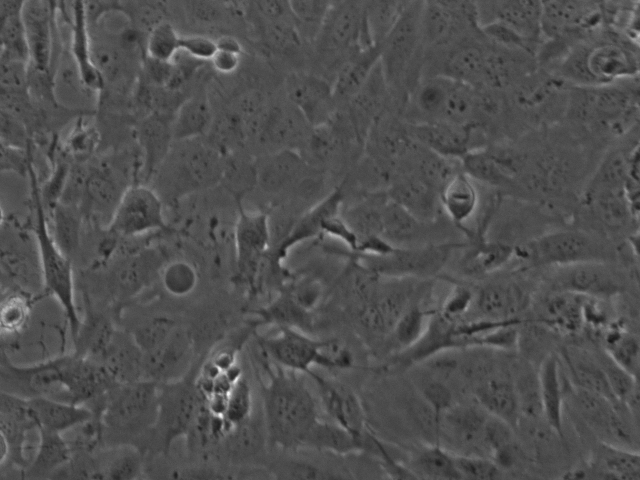  (c)  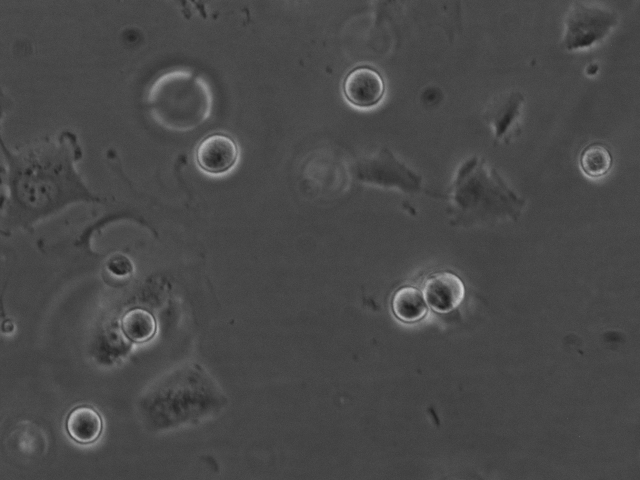  (e) | 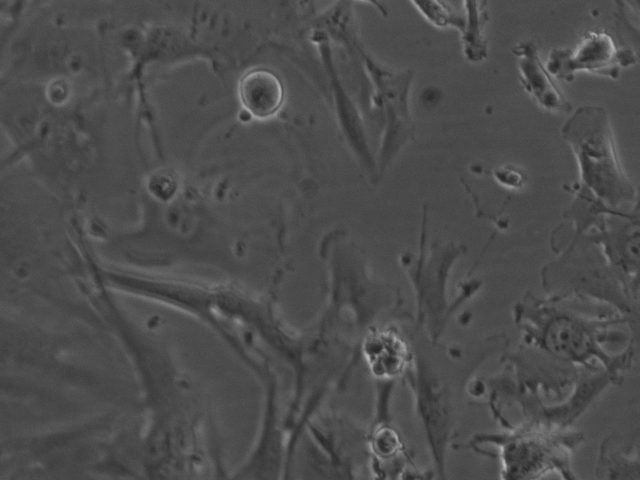  (**b**)  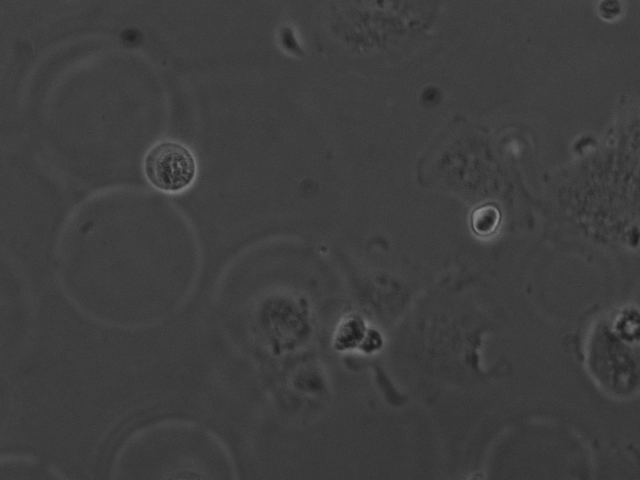  (d) |
| --- | --- |

**Figure S3.** Typical cell morphology 48 hours post radiotherapy and chemotherapy for experiments in Figure 1 and Figure S1. (a) T98G Ctl. (b) T98G + 20 Gy. (c) T98G + 2 Gy. (d) T98G + Pac. (e) T98G + Pac + 20 Gy. Scale bar is 100 μm.

| 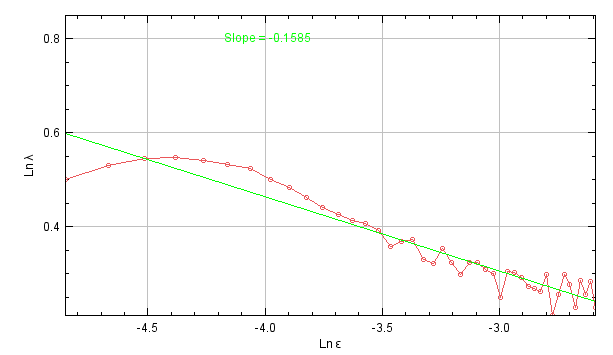  (**a**)  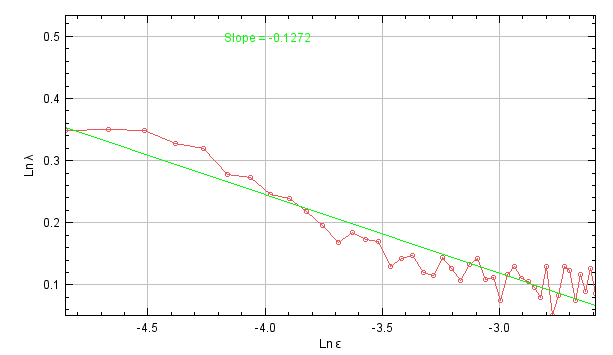  (c)  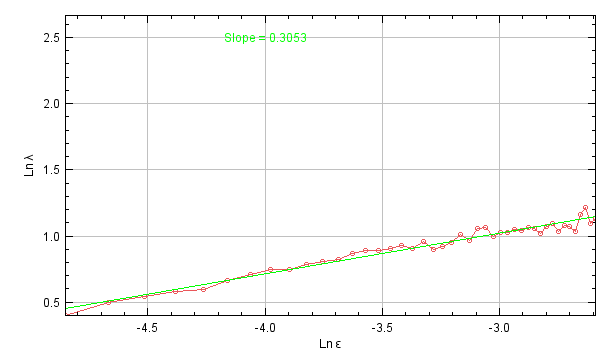  (e) | 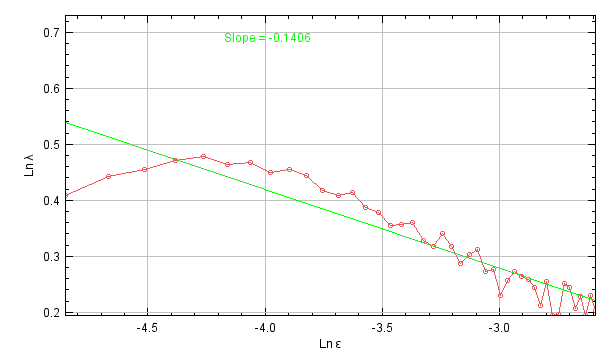  (**b**)  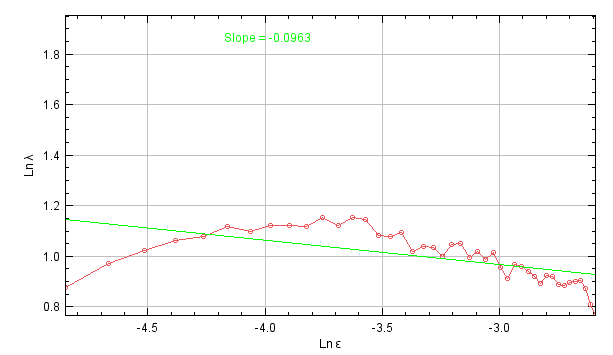  (d) |
| --- | --- |

**Figure S4.** Typical lacunarity-based cell morphometry 48 hours post radiotherapy and chemotherapy for experiments in Figure 1 and Figure S1. The logarithm of lacunarity (λ) was plotted against the logarithm of box size (ε). The closer the slope is to 0, the more homogeneous the image. (a) T98G Ctl. (b) T98G + 20 Gy. (c) T98G + 2 Gy. (d) T98G + Pac. (e) T98G + Pac + 20 Gy.

| 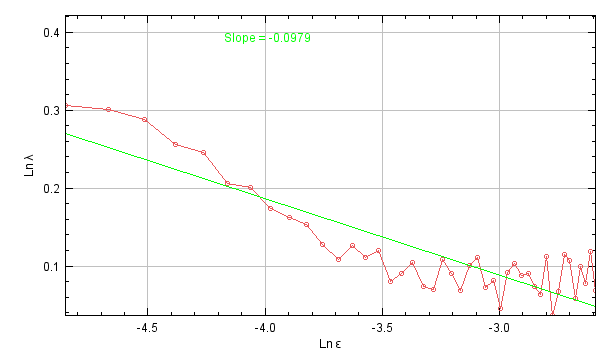  (**a**)  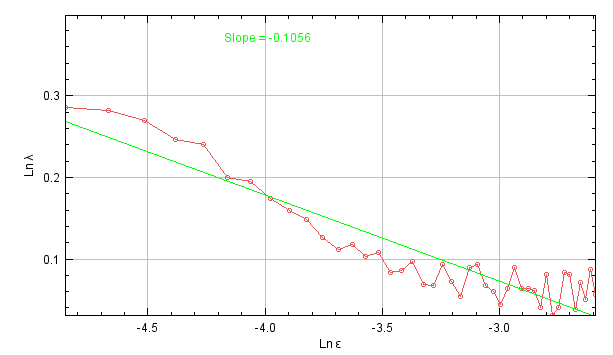  (c)  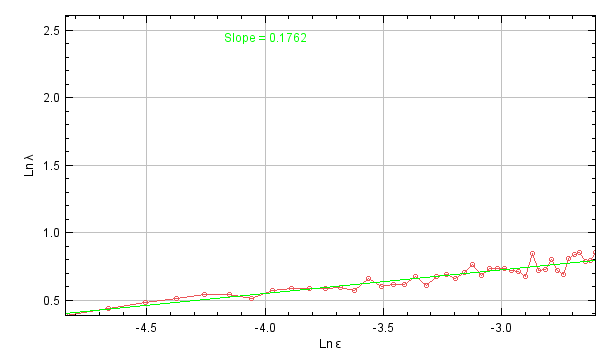  (e) | 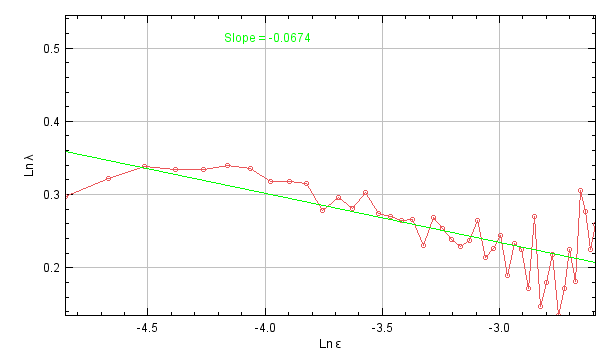  (**b**)  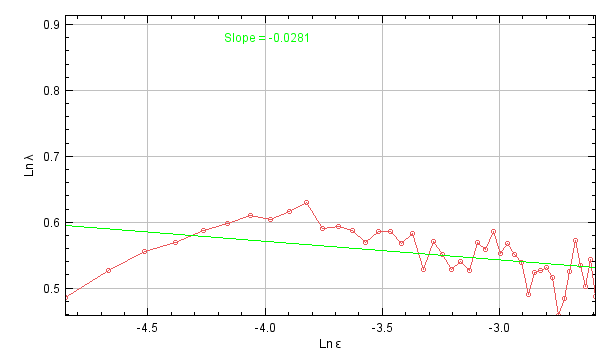  (d) |
| --- | --- |

**Figure S5.** Typical lacunarity-based cell morphometry 96 hours post radiotherapy and chemotherapy for experiments in Figure 1 and Figure S1. The logarithm of lacunarity (λ) was plotted against the logarithm of box size (ε). The closer the slope is to 0, the more homogeneous the image. (a) T98G Ctl. (b) T98G + 20 Gy. (c) T98G + 2 Gy. (d) T98G + Pac. (e) T98G + Pac + 20 Gy.
